# Multimodal Evidence for the Effectiveness of a Digital Behaviour Change Platform: Behavioural, Biomarker, and EEG Outcomes in University Students

**DOI:** 10.64898/2026.07.23.740387

**Authors:** Rae Dauphine, Mathew Hammerstrom, Olav E. Krigolson

## Abstract

Digital behaviour change interventions have emerged as a promising approach for improving health, well-being, and performance, yet relatively few studies have evaluated their effects using objective biological and neurophysiological measures. The present study examined the effectiveness of Autonomic, a neuroscience-informed digital coaching platform designed to improve student well-being through personalized behavioural interventions targeting sleep, stress, mood, energy, and focus. Thirty university students engaged with the platform for ten weeks and completed behavioural assessments, biomarker collection, and electroencephalographic (EEG) testing at baseline, five weeks, and ten weeks. Behavioural outcomes included self-reported ratings of focus, energy, sleep, mood, and stress. Biological measures included salivary cortisol and tear fluid dopamine concentrations. EEG assessments included resting-state recordings, frontal theta activity during a working memory task, and N200/P300 event-related potentials during a visual oddball task. Significant improvements were observed in self-reported focus, energy, and sleep quality across the intervention. Electrophysiological measures demonstrated reduced frontal theta power during working memory and shorter P300 latencies during attentional processing, consistent with more efficient cognitive processing following the intervention. Although cortisol, dopamine, and resting-state EEG measures did not reach statistical significance, all exhibited changes in the predicted direction. Collectively, these findings provide converging behavioural, biological, and neurophysiological evidence supporting the effectiveness of Autonomic. More broadly, the study demonstrates the value of combining objective biomarkers and EEG with traditional behavioural assessments when evaluating digital behaviour change interventions and highlights the potential of neuroscience-informed coaching platforms to improve health, well-being, and cognitive functioning in university students.

## Introduction

Durable improvements in health and performance require more than awareness of what should change; they require the successful regulation of repeated choices and actions. Sleep, physical activity, stress management, and self-care are expressed through everyday behaviour, making the capacity to alter established routines central to both health promotion and disease prevention (Michie et al., 2011). Behavioural interventions address this challenge through theoretically grounded methods that make target actions observable, manageable, and repeatable. Among the most established methods are self-monitoring, goal setting, performance feedback, prompts, and procedures intended to support habit development (Abraham & Michie, 2008; Michie et al., 2013). Their value lies in helping people convert broad intentions into specific actions, regulate progress when circumstances change, and gradually stabilize health-promoting routines (Spring et al., 2020). Across physical activity, diet, medication adherence, and psychological well-being, reviews indicate that interventions incorporating these techniques can produce meaningful change (Michie et al., 2011; Spring et al., 2020). This behavioural foundation is therefore central to digital health systems intended to influence wellness and human performance over time.

The university years provide a demanding context in which to test such an approach. Students must establish new routines while simultaneously adapting to greater academic responsibility, increased independence, financial strain, and substantial changes in their social and living environments (Arnett, 2000; Bewick et al., 2010). Under these conditions, sleep, exercise, nutrition, stress regulation, and mental health can deteriorate together rather than as independent problems. These patterns matter because less adaptive health behaviour during university is associated with greater psychological distress, poorer well-being, and weaker academic outcomes (Deliens et al., 2014; Richardson et al., 2012). The developmental timing is also important: emerging adulthood is a period in which behavioural patterns can consolidate and continue beyond graduation (Nelson et al., 2008). Supporting students as they construct their own routines may therefore yield immediate benefits while also shaping the longer-term behavioural repertoire carried into adult life.

Delivering behavioural support digitally may be especially well suited to this population. Smartphone-based systems can accompany users across settings and provide individualized monitoring, goals, and feedback at the moments when behaviour occurs, without requiring the staffing and scheduling demands of repeated face-to-face sessions (Murray et al., 2016; Webb et al., 2010). That delivery model is relevant in post-secondary institutions, where the need for student support can exceed the capacity of conventional services. Evidence synthesized across digital interventions shows benefits for outcomes that include physical activity, sleep, stress, and mental health (Firth et al., 2017; Webb et al., 2010). Digital delivery thus offers a practical route for extending structured behaviour-change support across a large student body while retaining responsiveness to the needs and progress of individual users.

Autonomic applies this model through a neuroscience-informed mobile platform created for university students. Its coaching addresses sleep, stress, mood, energy, and focus as interacting contributors to well-being and performance. Brief daily exchanges incorporate self-observation, individualized feedback, goal formation, and habit-building procedures so that support is integrated into users’ routines rather than confined to periodic appointments. The platform is intended to combine individualized guidance with the reach and low resource burden of mobile delivery. However, a theoretically informed design does not by itself establish that engagement produces meaningful change. Evaluation should determine whether changes appear not only in users’ ratings but also in biological and neural processes relevant to stress regulation and cognition. The present study therefore examined university students with self-report, biomarker, and electroencephalographic (EEG) measures collected across their engagement with Autonomic.

Given that Autonomic targets multiple aspects of well-being, including stress management, emotional regulation, energy, focus, and healthy lifestyle behaviors, its effectiveness is most appropriately evaluated using a multimodal assessment framework. Cortisol was included as a biological marker of hypothalamic-pituitary-adrenal (HPA) axis activity and physiological stress, as chronic stress and poor self-regulation are associated with elevated cortisol levels and dysregulated stress responses (Kudielka et al., 2009; McEwen, 1998). Dopamine was assessed because dopaminergic systems play a central role in motivation, reward processing, goal-directed behavior, and learning, all of which are relevant to successful behavior change (Schultz, 2016; Wise, 2004). Resting-state electroencephalography (EEG) was included to provide an objective measure of baseline neural function, as patterns of ongoing neural activity have been linked to cognitive performance, arousal, and psychological well-being (Başar, 2012; Klimesch, 1999). In addition, frontal theta power measured during an n-back working memory task was used as an index of cognitive engagement and resource allocation, with greater frontal theta activity consistently associated with the recruitment of cognitive control processes during demanding cognitive tasks (Cavanagh & Frank, 2014; Mitchell et al., 2008). Finally, the N200 and P300 event-related potentials elicited during an oddball task were included because they provide established neural indices of attentional allocation, stimulus evaluation, and cognitive control, processes that are fundamental to effective self-regulation and adaptive behavior (Folstein & Van Petten, 2008; Polich, 2007). Collectively, these measures provide complementary biological and neurocognitive indicators through which changes associated with engagement in the Autonomic platform may be assessed.

The primary objective of the present study was to evaluate the effectiveness of the Autonomic platform in promoting improvements in psychological well-being, biological function, and neural indicators of cognitive performance in university students. To accomplish this, participants completed self-report assessments and provided biological and EEG measures at baseline, following five weeks of platform use, and again after ten weeks of engagement with Autonomic. Based on the extensive literature supporting behavior change interventions and digital health technologies, it was hypothesized that participants would demonstrate progressive improvements in self-reported well-being across the intervention period. Consistent with these anticipated improvements, we further predicted that biological markers associated with stress and adaptive functioning would exhibit favorable changes over time, including reductions in cortisol and alterations in dopamine concentrations. At the neural level, we hypothesized that engagement with the platform would be associated with changes in resting-state EEG activity indicative of improved cognitive state, enhanced frontal theta activity during working memory performance reflecting more effective cognitive engagement, and modulation of the N200 and P300 event-related potentials during an oddball task consistent with improvements in attentional processing and cognitive control. Collectively, these predicted changes would provide converging evidence that sustained engagement with the Autonomic platform is associated with measurable improvements in psychological, biological, and neural indices relevant to health, well-being, and performance.

## Methods

### Participants

Participants were recruited from the general undergraduate student population at the University of Victoria. Thirty individuals with no known neurological impairments and normal or corrected-to-normal vision took part in the experiment. The final sample was evenly split between female and male participants: 15 females (50%) and 15 males (50%). Age was not collected. All 30 participants completed the three assessment sessions and contributed usable data to every reported behavioural, biomarker, and EEG analysis. Participants received CAD $100 and provided written informed consent. The study was approved by the University of Victoria Human Research Ethics Board (protocol 25-0254) and conducted in accordance with the ethical standards prescribed in the 1964 Declaration of Helsinki and its subsequent amendments.

### Experimental Design

Participants completed a ten-week experimental protocol during which they used the Autonomic web-based application on a daily basis (https://www.getautonomic.com/). In brief, Autonomic is a digital health and performance application designed to support student well-being through brief, personalized, neuroscience-informed coaching. The application uses daily check-ins and micro-coaching sessions to help users monitor and improve five core domains related to cognitive and emotional functioning: sleep quality, stress, mood, energy, and focus. Users receive individualized guidance intended to promote habit formation, self-regulation, and improved day-to-day functioning, with sessions designed to be completed in approximately three minutes.

Before beginning the Autonomic program (baseline), after five weeks of platform use (midpoint), and after ten weeks of platform use (endpoint), participants attended the laboratory and completed the five Autonomic ratings, EEG testing, and biomarker collection for cortisol and dopamine. The primary analyses therefore compared three time points: baseline, midpoint, and endpoint.

### Biomarker Measurement and Analysis

Testing was conducted in a quiet, well-lit laboratory. Participants were asked not to eat or drink for at least one hour before sample collection. Saliva was collected using a Salivette cotton swab (Sarstedt, catalogue no. 51.1534), which participants held in the mouth for two minutes. Tear fluid was collected using a Clement Clarke Schirmer tear-test strip placed in the eye for five minutes or until saturated. The strip was then cut with sterile scissors into 300 µL of phosphate-buffer solution and agitated for 60 minutes. Samples were initially stored at −20 °C and transferred within one week to −80 °C storage in the University of Victoria Health Core Laboratory. Collection time was not standardized across visits because testing depended on laboratory and participant availability. Salivary cortisol is a well-established non-invasive measure of biologically active free cortisol and is commonly used as an index of hypothalamic-pituitary-adrenal (HPA) axis activity (Inder & Dimeski, 2012). Tear-fluid dopamine can also be detected and quantified in humans (Sharma et al., 2019). Cortisol was quantified using the Human Cortisol Competitive ELISA Kit (Thermo Fisher Scientific, catalogue no. EIAHCOR), and dopamine was quantified using the Dopamine Competitive ELISA Kit (Thermo Fisher Scientific, catalogue no. EEL144). Manufacturer-reported intra-and inter-assay coefficients of variation were below 10%. The assays were performed by a trained research technician in the University of Victoria Health Core Laboratory.

### EEG Assessment and Data Acquisition

Following biomarker collection, participants were fitted with the X.on wireless EEG system (Brain Products GmbH, Gilching, Germany). Seven passive sponge-based Ag/AgCl electrodes were positioned at F3, F4, C3, Cz, C4, P3, and P4 according to the international 10-20 system; the reference and ground electrodes were contained in an ear clip. EEG was sampled at 500 Hz and transmitted wirelessly via Bluetooth to PEER (Version 4.2; BrainWave Software Inc., Victoria, BC), which was used for stimulus presentation, event marking, and EEG acquisition.

Participants completed three experimental protocols – a steady state eyes-open/eyes-closed assessment (three minutes in each condition), a standard two-back nBack task (4 blocks of 50 trials), and a standard visual oddball task (4 blocks of 60 trials). All of the experimental tasks were done using the aforementioned PEER software (www.peereeg.com).

### Data Processing and Analysis

#### Behavioural Data

At each time point, participants completed the same five brief Autonomic check-in items: “Rate your focus level,” “Rate your energy level,” “Rate your sleep quality last night,” “Rate your mood level,” and “Rate your stress level.” Focus was defined as the ability to work without distraction; energy as the energy available for thinking, focusing, and getting things done; sleep quality as how restful and uninterrupted the previous night’s sleep was and how refreshed the participant felt upon waking; mood as the participant’s overall state of mind or feeling; and stress as how overwhelmed or under pressure the participant felt. Responses were recorded on 1-10 sliders labelled Low, Medium, and High and were multiplied by 10 for analysis and presentation. Higher scores indicated greater focus, energy, sleep quality, mood, or stress, respectively.

#### Electroencephalographic Data

All EEG processing was conducted in MATLAB using the EEGLAB analysis package and custom scripts generated by the senior author (www.github.com/krigolson). First, data were downsampled to 250 Hz. Next, we applied a dual-pass phase-free Butterworth filter with a band-pass of 0.5 to 30 Hz. The processing of the EEG data at this point became task dependent.

#### Steady State

Eyes-open and eyes-closed blocks were identified from event markers, with two seconds trimmed from the beginning and end of each block to avoid startup and end noise. The data were then segmented into 1-second epochs with 0.5-second overlap, and epochs were rejected if any scalp channel exceeded ±100 µV. Power spectral density was then computed for each clean epoch, averaged across clean epochs, and averaged across F3 and F4 and P3 and P4 to create anterior and posterior regions of interest. Spectra were summarized in centered 1-Hz bins from 1 to 30 Hz separately for eyes-open and eyes-closed conditions at each session. Band power was computed for the delta (1 to 3 Hz), theta (4 to 7 Hz), alpha (8 to 12 Hz), and beta (13 to 30 Hz) bands for both the anterior and posterior regions of interest for each condition (eyes-open and eyes-closed) and time point (baseline, midpoint, endpoint) for each participant.

Alpha reactivity was quantified as the difference in posterior alpha power between the eyes-closed and eyes-open conditions. Specifically, alpha power was computed over the 8 to 12 Hz band at the posterior region of interest, log10 transformed, and then calculated as eyes-closed minus eyes-open log10 alpha power. Larger values therefore indicate greater alpha enhancement during eye closure relative to eye opening, whereas smaller values indicate reduced eyes-open/eyes-closed alpha differentiation. As with band power, alpha reactivity was computed for each time point (baseline, midpoint, endpoint) for each participant.

#### nBack

For the n-back theta analysis, task data were segmented into 1000 ms epochs with 500 ms overlap, and epochs were rejected if they contained all-zero signals or if any filtered scalp channel exceeded ±150 µV. Power spectral density was computed for clean epochs, theta power was quantified from 4 to 7 Hz, and relative theta was calculated as theta power divided by total power from 0.5 to 30 Hz. The primary frontal outcome was log10 relative theta power averaged across F3, F4, and Cz for each time point (baseline, midpoint, endpoint) and participant.

#### Oddball

Standard and Target trials were identified from the visual oddball event markers, and ERP epochs were extracted from −200 to 600 ms relative to stimulus onset. Epochs were baseline-corrected using the −200 to 0 ms pre-stimulus interval and rejected if the peak-to-peak voltage difference exceeded 150 µV on any scalp channel. Participant-level ERPs were averaged separately for Standard and Target trials at each session, and difference waveforms were computed. The N200 was quantified from an F3/F4/Cz region of interest, and the P300 was quantified from a P3/P4 region of interest. Using the difference waveforms, N200 latency was identified for each participant as the most negative peak between 150 and 300 ms, and P300 latency was identified as the most positive peak between 300 and 600 ms; if the detected peak occurred at the edge of the search window, the best within-range local peak was used instead. Peak amplitude was then quantified as the mean voltage within ±20 ms of the identified peak latency.

### Statistical Analysis

Each dependent variable was submitted to a one-factor repeated-measures analysis of variance with three levels of time (baseline, midpoint, endpoint). Planned linear contrasts assessed systematic change across the three assessments (Maxwell et al., 2018; Rosenthal et al., 2000). All statistical assumptions were tested and met. An alpha level of .05 was used for all tests. Cohen’s d or partial eta squared (ηp²) is reported as the effect-size estimate, as appropriate. Error bars in all figures represent 95% within-subject confidence intervals (Masson & Loftus, 2003). Analyses were conducted using JASP.

## Results

### Behavioural

A series of repeated-measures ANOVAs were conducted to examine changes across the three assessment points (baseline, midpoint, endpoint) for energy, focus, mood, sleep quality, and stress (see Figure 1). For focus, there was a significant main effect of time, F(2, 58) = 6.189, p = .004, ηp² = .176. The planned linear trend was also significant, F(1, 29) = 7.746, p = .009, ηp² = .211, reflecting a 24.78% increase in focus over time points. A similar result was found for energy: a main effect of time, F(2, 58) = 3.943, p = .025, ηp² = .120, and a 19.32% increase in energy ratings. The planned linear trend for energy was also significant, F(1, 29) = 7.104, p = .012, partial η² = .197. For sleep, the main effect of time approached significance, F(2, 58) = 2.964, p = .059, ηp² = .093. The planned linear trend was significant, F(1, 29) = 6.416, p = .017, partial η² = .181, reflecting a 15.39% increase in sleep ratings. No significant effects of time were observed for mood or stress, ps > .05. However, descriptively, mood increased by 8.05%, and stress decreased by 10.14% from baseline to endpoint (see Figure 1).

**Figure 1.**
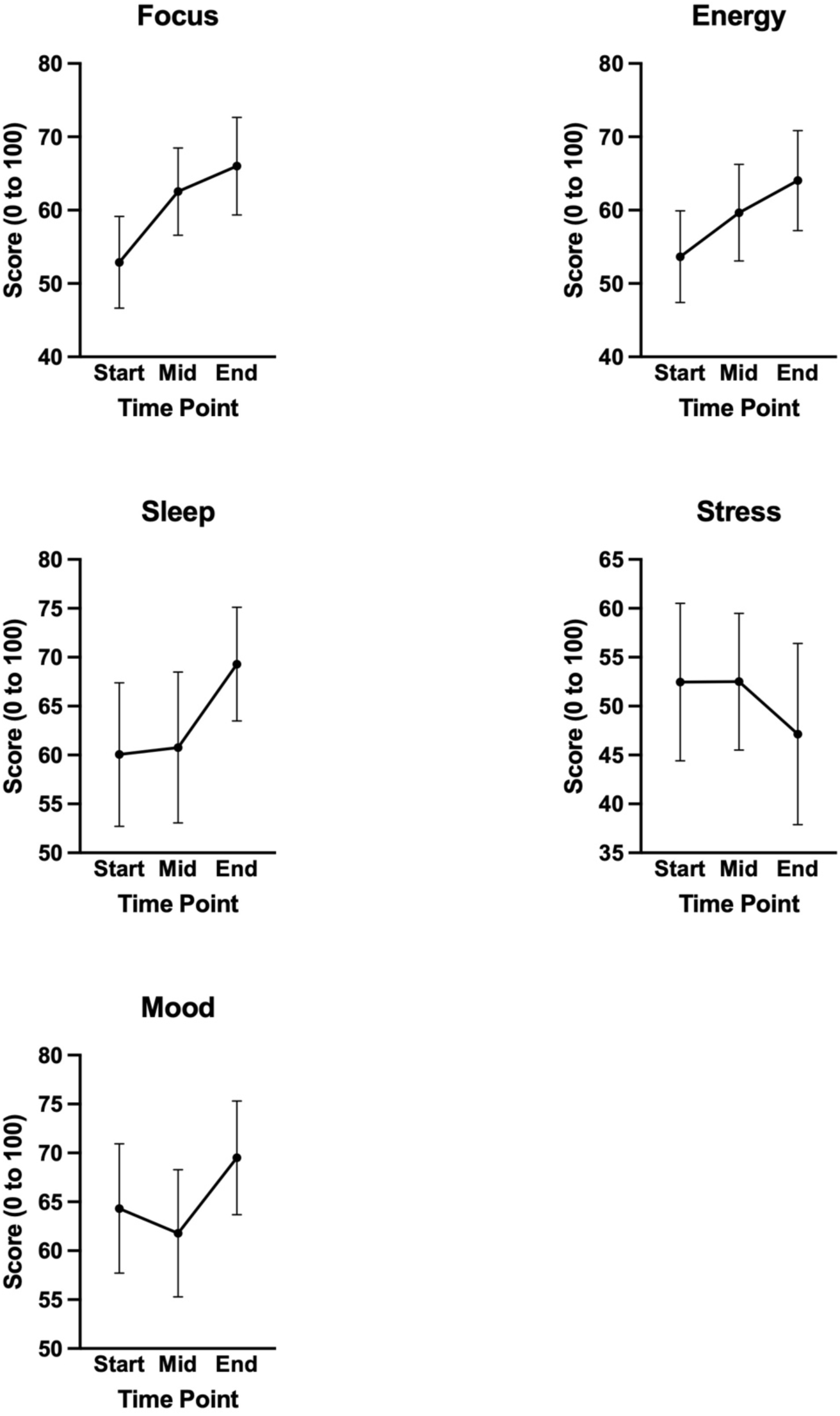
Ratings for focus, energy, sleep, stress, and mood at the three experimental time points: baseline, midpoint, and endpoint.

### Biomarker Data

#### Cortisol

A repeated-measures ANOVA (baseline, midpoint, endpoint) showed no significant effect of time, F(2, 58) = 1.11, p = .336, ηp² = .037. The planned linear trend also was not significant, F(1, 29) = 1.10, p = .303, Cohen’s d = −0.19. However, we observed a 23.6% reduction in cortisol levels from baseline to endpoint (see Figure 2).

**Figure 2.**
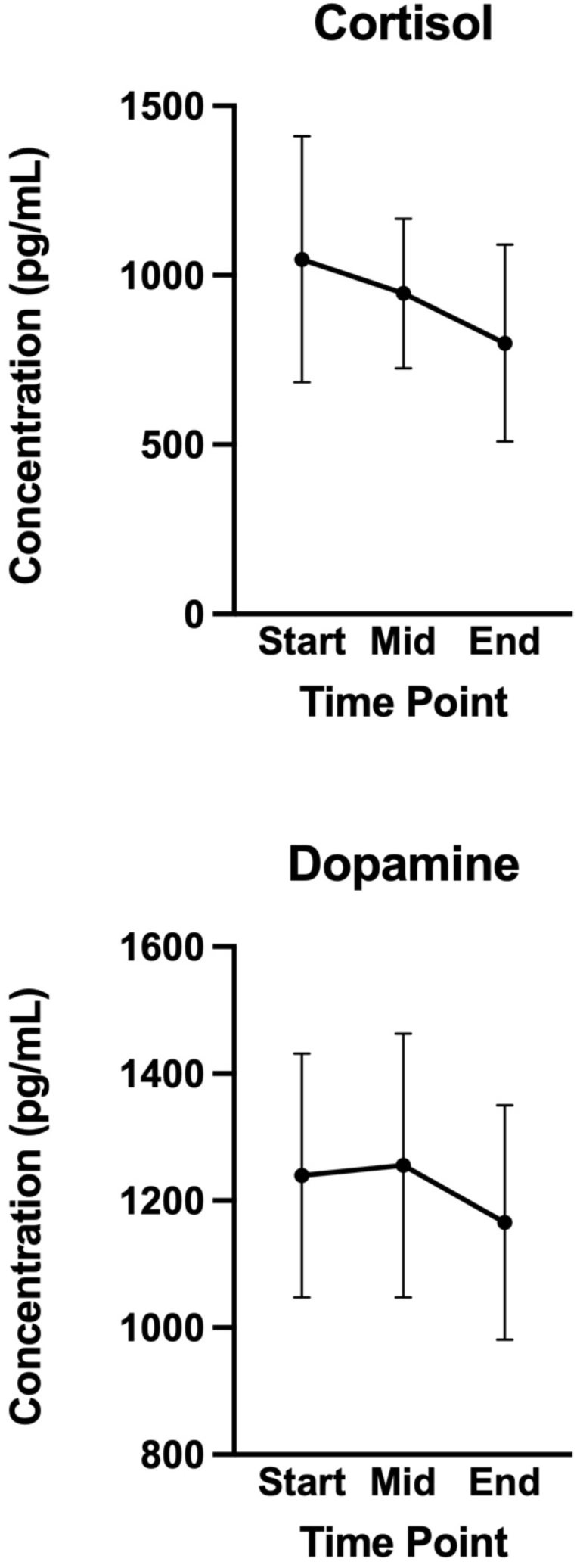
Changes in cortisol and dopamine levels as a function of experimental time point.

#### Dopamine

A repeated-measures ANOVA showed no significant effect of time point, F(2, 58) = 1.70, p = .192, ηp² = .055. The planned linear trend was also not significant, F(1, 29) = 1.46, p = .237, Cohen’s d = −0.22, with dopamine levels decreasing by 6.0% from baseline to endpoint (see Figure 2).

### Electroencephalographic Data

#### Steady State

As expected, across the experiment, alpha power was substantially greater during eyes closed than eyes open, producing a main effect of condition, F(1, 29) = 56.74, p < .001, ηp² = .662 (see Figure 3). Over the three experimental time points, there was no effect, F(2, 58) = 2.54, p = .088, ηp² = .081. However, the planned linear trend showed a 20.5% decrease across time points that approached significance, F(1, 29) = 2.86, p = .101, Cohen’s d = −0.31. None of the band power frequency bands that were analyzed showed any temporal effects, ps > .05. For EO/EC alpha reactivity, the repeated-measures ANOVA revealed a medium-sized trend for time, F(2, 58) = 2.54, p = .088, ηp² = .081. The planned linear trend showed a 20.5% decrease across time points, F(1, 29) = 2.86, p = .101, Cohen’s d = −0.31.

**Figure 3.**
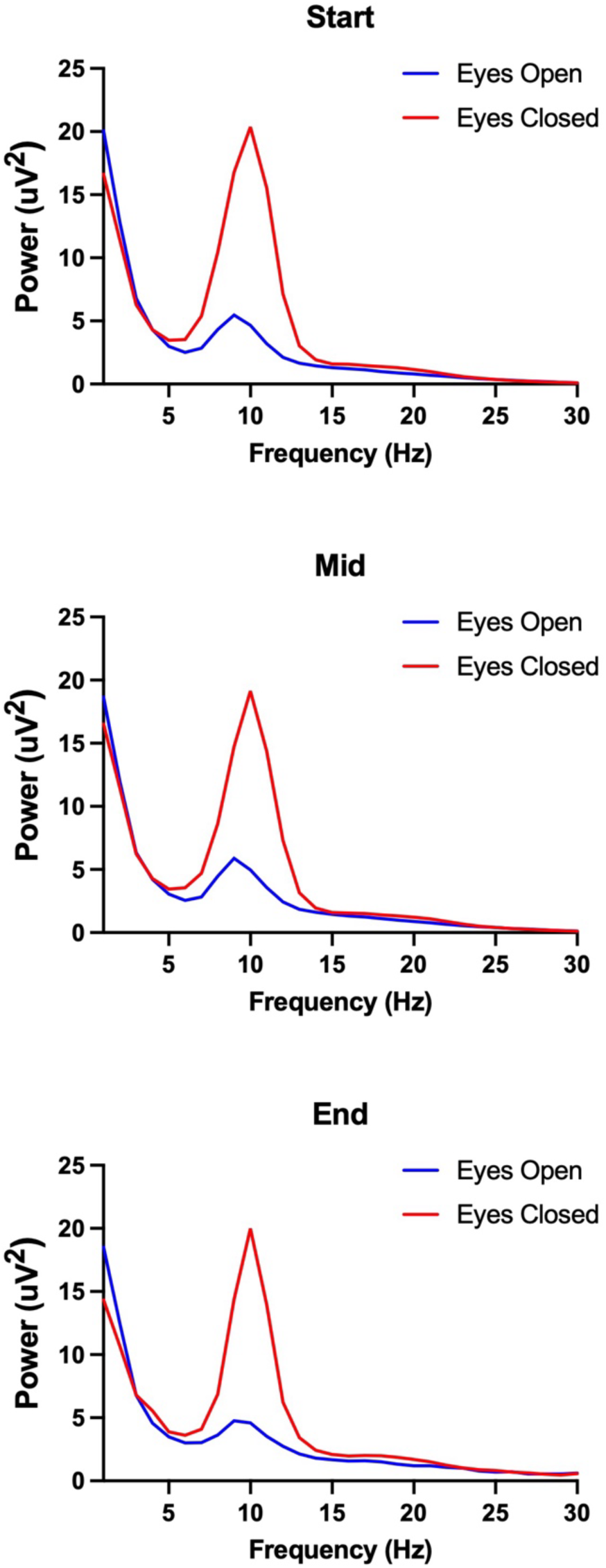
EEG power spectra for the eyes-open and eyes-closed conditions at baseline, midpoint, and endpoint.

**Figure 4.**
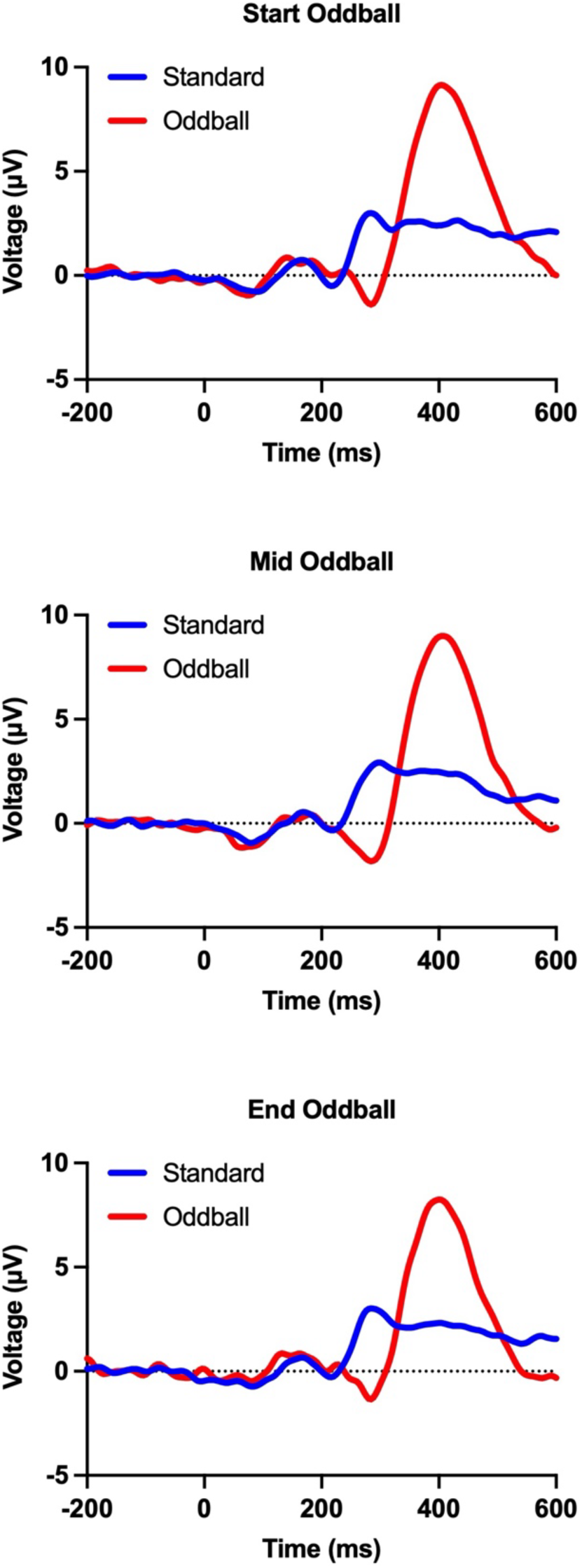
Grand average ERP waveforms from the oddball task at the three time points: baseline, midpoint, and endpoint.

#### nBack

The analysis of relative frontal theta power showed an effect of time point, F(2, 58) = 6.60, p = .003, ηp² = .185, with a 7.9% decrease across time points, F(1, 29) = 6.34, p = .018, Cohen’s d = −0.46 (see Figure 5).

**Figure 5.**
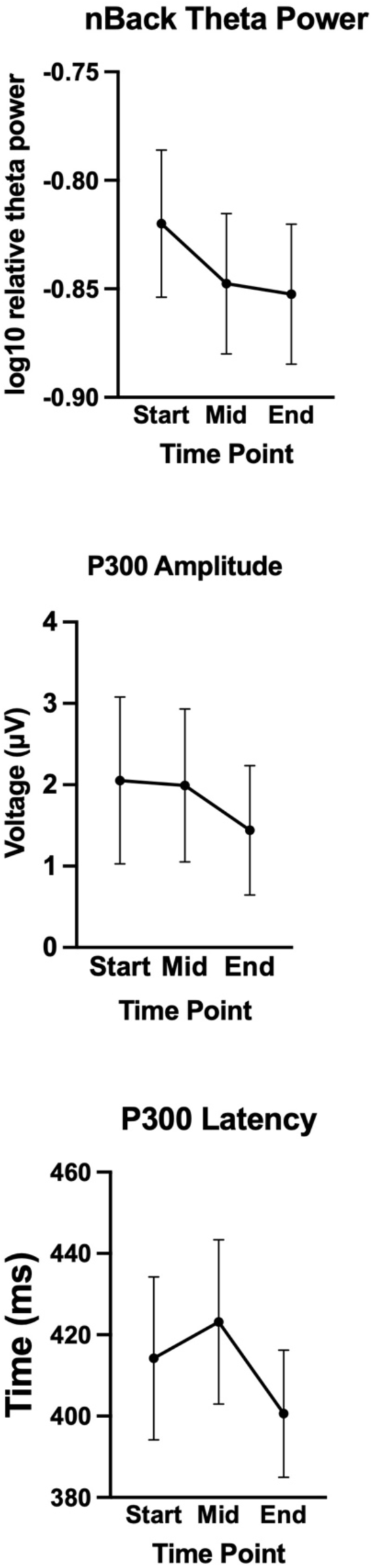
Medial frontal theta power, P300 amplitude, and P300 latency as a function of experimental time point (baseline, midpoint, endpoint).

#### Oddball

*N200.* No reliable N200 effects were observed; Target-minus-Standard N200 amplitude did not significantly vary across time points, F(2, 58) = 2.13, p = .129, ηp² = .068, and the planned linear trend was not significant, F(1, 29) = 2.65, p = .114, Cohen’s d = −0.30.

*P300.* For the Target-minus-Standard P3 latency at P3/P4, there was a significant effect of time point, F(2, 58) = 3.48, p = .037, ηp² = .107; the planned linear trend was also significant, F(1, 29) = 4.84, p = .036, Cohen’s d = −0.40, reflecting a 13.6 ms, or 3.3%, decrease from baseline to endpoint. P3 amplitude at P3/P4 showed a non-significant effect of time point, F(2, 58) = 2.26, p = .114, ηp² = .072; the planned linear trend was trend-level, F(1, 29) = 3.14, p = .087, Cohen’s d = −0.32, reflecting a 29.9% decrease from baseline to endpoint (see Figure 5).

## Discussion

This study tested Autonomic across three levels of measurement: participants’ ratings of daily functioning, biological indices, and electrophysiological measures obtained during rest and cognitive performance. Across ten weeks, focus, energy, and sleep quality improved progressively, extending evidence that digitally delivered coaching can support well-being and self-regulation through individualized goals, feedback, and repeated self-observation (Michie et al., 2011; Murray et al., 2016; Spring et al., 2020). The self-report pattern was accompanied by lower frontal theta power during working memory and shorter P300 latency during attentional processing. These measures provide distinct information: frontal theta indexes the recruitment of cognitive control resources (Cavanagh & Frank, 2014; Mitchell et al., 2008), whereas P300 latency reflects the speed of stimulus evaluation and attentional processing (Polich, 2007). Cortisol, dopamine, and resting EEG did not change significantly, although each moved in the hypothesized direction. The contribution of the present results is therefore their convergence: Autonomic-related change was visible in subjective experience and in task-evoked brain activity, with supporting directional patterns in the biomarker data. Such triangulation addresses the need to evaluate digital health technologies across complementary levels of evidence rather than through self-report alone (Murray et al., 2016).

At the behavioural level, the clearest change was a 25% increase in focus, the largest self-reported effect in the study. Attention and concentration are directly relevant to academic success and can be undermined when students face inadequate sleep, elevated stress, and simultaneous demands (Bewick et al., 2010; Richardson et al., 2012). Energy and sleep quality also improved significantly, while mood rose and perceived stress declined in the predicted direction. This breadth is consistent with evidence that self-monitoring, individualized feedback, explicit goals, and habit-oriented methods can alter health behaviour and psychological well-being (Abraham & Michie, 2008; Michie et al., 2011, 2013; Webb et al., 2010). The findings are particularly relevant in the university context, where the transition into post-secondary study commonly disrupts routines and is accompanied by poorer sleep, greater stress, and diminished well-being (Arnett, 2000; Deliens et al., 2014). Autonomic treats these outcomes as a connected system rather than as isolated targets. The observed pattern supports the proposition that a multidomain, neuroscience-informed coaching platform can influence several behaviours relevant to academic functioning and long-term health during a period when enduring habits are still being formed (Nelson et al., 2008; Spring et al., 2020).

The electrophysiological findings provide important objective support for the behavioural improvements associated with Autonomic. Although resting-state EEG measures remained largely unchanged across the intervention, significant changes were observed during tasks that required active cognitive engagement. Specifically, participants demonstrated reduced frontal theta power during the n-back working memory task together with faster P300 latencies during the visual oddball task. Frontal midline theta is widely regarded as an index of cognitive control and the allocation of mental resources, with greater theta power generally reflecting increased recruitment of executive control processes during demanding cognitive tasks (Cavanagh & Frank, 2014; Mitchell et al., 2008). Likewise, the P300 component has long been recognized as a neural marker of attentional allocation and stimulus evaluation, with shorter latencies indicating faster classification and processing of task-relevant information (Polich, 2007). Considered together, these findings suggest that following ten weeks of engagement with Autonomic, participants were able to perform cognitive tasks while recruiting fewer neural resources and processing relevant information more rapidly. This pattern is consistent with the theory of neural efficiency, which proposes that improvements in cognitive performance are often accompanied by reductions in task-related neural activity as the brain becomes more efficient at allocating computational resources (Neubauer & Fink, 2009). Importantly, the absence of substantial changes in resting-state EEG further suggests that Autonomic does not fundamentally alter baseline brain activity, but rather improves the efficiency with which cognitive control networks are recruited when cognitive demands arise. These objective neural changes closely parallel the observed improvements in self-reported focus, providing converging evidence that behavioural gains associated with Autonomic are accompanied by measurable changes in brain function.

Although neither cortisol nor dopamine concentrations changed significantly across the intervention, both biomarkers exhibited changes in the predicted direction. Cortisol levels decreased by approximately 24% over the ten-week intervention, consistent with reduced physiological stress and improved regulation of the hypothalamic-pituitary-adrenal (HPA) axis (Kudielka et al., 2009; McEwen, 1998). Dopamine concentrations also showed a modest reduction. Although this finding should be interpreted cautiously, it may reflect reduced reliance on dopaminergic reward prediction error signalling as healthy behaviours become increasingly learned and habitual over the course of the intervention. Dopamine plays a central role in reinforcing behavioural adaptations by signalling discrepancies between expected and experienced outcomes, with this signal diminishing as behaviours become predictable and well learned (Schultz, 2016). Consequently, a reduction in dopamine following sustained engagement with Autonomic may be consistent with participants transitioning from effortful behaviour change toward more automatic and stable behavioural patterns. Given the relatively small sample size and substantial inter-individual variability associated with biological markers, these findings should be considered preliminary. Nevertheless, the directional changes observed in both biomarkers complement the behavioural and electrophysiological findings and suggest that sustained engagement with Autonomic may influence not only psychological functioning but also the biological mechanisms that support adaptive behaviour.

Viewed as a whole, the study yielded a coherent pattern across behavioural, biological, and neurophysiological observations. Better focus, energy, and sleep coincided with task-evoked EEG changes consistent with more efficient processing, while the biomarker trajectories were compatible with reduced physiological stress and adaptive regulation. The fact that some outcomes did not reach statistical significance does not erase the consistency in direction across levels of analysis. Using surveys, biochemical measures, and EEG also allowed the platform to be examined from complementary perspectives rather than judged solely by what participants reported. As digital health tools become more common, combining subjective outcomes with physiological and neural measures can provide a fuller test of intervention efficacy (Murray et al., 2016). On that basis, the current findings indicate that neuroscience-informed digital coaching may be associated with changes spanning everyday experience, biological regulation, and brain function.

The implications extend to human performance because reliable performance depends on the state of the systems that support it. Attention, working memory, and cognitive control are vulnerable to both stress and insufficient sleep, while frontal theta and the P300 capture processes involved in control, resource allocation, and the evaluation of task-relevant information (Cavanagh & Frank, 2014; Lim & Dinges, 2010; Lupien et al., 2009). Autonomic may therefore contribute to performance by improving the conditions under which cognition operates: lower stress, better sleep, greater energy and focus, and more efficient task-related neural processing. In this sense, the platform’s potential value is not limited to subjective wellness but includes readiness for sustained and effective action across study, work, and daily life. Direct assessment of behavioural performance will be necessary to determine whether this multimodal pattern produces durable gains on objective tasks and consequential real-world outcomes.

In conclusion, the present study provides encouraging evidence that the Autonomic digital coaching platform can produce measurable improvements in student well-being that extend beyond self-reported behaviour to include objective neurophysiological and biological indicators of cognitive function. By integrating behavioural science with multimodal physiological assessment, this work demonstrates a promising approach for evaluating digital health interventions and highlights the potential of neuroscience-informed coaching to promote healthier, more adaptive patterns of behaviour. As objective measures become increasingly accessible through advances in wearable technologies and digital health platforms, approaches such as those described here may play an important role in the future development, validation, and personalization of behaviour change interventions.

## Data Availability

Participant-level data are not publicly available because of privacy and commercial restrictions.

## References

Abraham, C., & Michie, S. (2008). A taxonomy of behavior change techniques used in interventions. Health Psychology, 27(3), 379–387. 10.1037/0278-6133.27.3.379

Arnett, J. J. (2000). Emerging adulthood: A theory of development from the late teens through the twenties. American Psychologist, 55(5), 469–480. 10.1037/0003-066X.55.5.469

Başar, E. (2012). A review of alpha activity in integrative brain function: Fundamental physiology, sensory coding, cognition and pathology. International Journal of Psychophysiology, 86(1), 1–24. 10.1016/j.ijpsycho.2012.07.002

Bewick, B., Koutsopoulou, G., Miles, J., Slaa, E., & Barkham, M. (2010). Changes in undergraduate students’ psychological well-being as they progress through university. Studies in Higher Education, 35(6), 633–645. 10.1080/03075070903216643

Cavanagh, J. F., & Frank, M. J. (2014). Frontal theta as a mechanism for cognitive control. Trends in Cognitive Sciences, 18(8), 414–421. 10.1016/j.tics.2014.04.012

Deliens, T., Deforche, B., De Bourdeaudhuij, I., & Clarys, P. (2014). Determinants of eating behaviour in university students: A qualitative study using focus group discussions. BMC Public Health, 14, 53. 10.1186/1471-2458-14-53

Firth, J., Torous, J., Nicholas, J., Carney, R., Rosenbaum, S., & Sarris, J. (2017). The efficacy of smartphone-based mental health interventions for depressive symptoms: A meta-analysis of randomized controlled trials. World Psychiatry, 16(3), 287–298. 10.1002/wps.20472

Folstein, J. R., & Van Petten, C. (2008). Influence of cognitive control and mismatch on the N2 component of the ERP: A review. Psychophysiology, 45(1), 152–170. 10.1111/j.1469-8986.2007.00602.x

Grabner, R. H., Stern, E., & Neubauer, A. C. (2006). Neural efficiency and intelligence: Insights from neuroimaging studies. In P. A. Vernon (Ed.), The Cambridge handbook of intelligence (pp. 101–119). Cambridge University Press.

Inder, W. J., & Dimeski, G. (2012). *Measurement of salivary cortisol in* 2012: Laboratory *techniques and clinical indications*. Clinical Endocrinology, 77(5), 645–651.

Klimesch, W. (1999). EEG alpha and theta oscillations reflect cognitive and memory performance: A review and analysis. Brain Research Reviews, 29(2–3), 169–195. 10.1016/S0165-0173(98)00056-3

Kudielka, B. M., Hellhammer, D. H., & Wüst, S. (2009). Why do we respond so differently? Reviewing determinants of human salivary cortisol responses to challenge. Psychoneuroendocrinology, 34(1), 2–18. 10.1016/j.psyneuen.2008.10.004

Lim, J., & Dinges, D. F. (2010). A meta-analysis of the impact of short-term sleep deprivation on cognitive variables. Psychological Bulletin, 136(3), 375–389. 10.1037/a0018883

Lupien, S. J., McEwen, B. S., Gunnar, M. R., & Heim, C. (2009). Effects of stress throughout the lifespan on the brain, behaviour and cognition. Nature Reviews Neuroscience, 10(6), 434– 445. 10.1038/nrn2639

Masson, M. E. J., & Loftus, G. R. (2003). Using confidence intervals for graphically based data interpretation. Canadian Journal of Experimental Psychology/Revue canadienne de psychologie expérimentale, 57(3), 203–220. 10.1037/h0087426

Maxwell, S. E., Delaney, H. D., & Kelley, K. (2018). Designing experiments and analyzing data: A model comparison perspective (3rd ed.). Routledge.

McEwen, B. S. (1998). Protective and damaging effects of stress mediators. New England Journal of Medicine, 338(3), 171–179. 10.1056/NEJM199801153380307

Michie, S., Richardson, M., Johnston, M., Abraham, C., Francis, J., Hardeman, W., Eccles, M. P., Cane, J., & Wood, C. E. (2013). The behavior change technique taxonomy (v1) of 93 hierarchically clustered techniques: Building an international consensus for the reporting of behavior change interventions. Annals of Behavioral Medicine, 46(1), 81–95. 10.1007/s12160-013-9486-6

Michie, S., van Stralen, M. M., & West, R. (2011). The behaviour change wheel: A new method for characterising and designing behaviour change interventions. Implementation Science, 6, 42. 10.1186/1748-5908-6-42

Mitchell, D. J., McNaughton, N., Flanagan, D., & Kirk, I. J. (2008). Frontal-midline theta from the perspective of hippocampal “theta.” Progress in Neurobiology, 86(3), 156–185. 10.1016/j.pneurobio.2008.09.005

Murray, E., Hekler, E. B., Andersson, G., Collins, L. M., Doherty, A., Hollis, C., Rivera, D. E., West, R., & Wyatt, J. C. (2016). Evaluating digital health interventions: Key questions and approaches. American Journal of Preventive Medicine, 51(5), 843–851. 10.1016/j.amepre.2016.06.008

Nelson, M. C., Story, M., Larson, N. I., Neumark-Sztainer, D., & Lytle, L. A. (2008). Emerging adulthood and college-aged youth: An overlooked age for weight-related behavior change. Obesity, 16(10), 2205–2211. 10.1038/oby.2008.365

Neubauer, A. C., & Fink, A. (2009). Intelligence and neural efficiency. Neuroscience & Biobehavioral Reviews, 33(7), 1004–1023. 10.1016/j.neubiorev.2009.04.001

Polich, J. (2007). Updating P300: An integrative theory of P3a and P3b. Clinical Neurophysiology, 118(10), 2128–2148. 10.1016/j.clinph.2007.04.019

Richardson, M., Abraham, C., & Bond, R. (2012). Psychological correlates of university students’ academic performance: A systematic review and meta-analysis. Psychological Bulletin, 138(2), 353–387. 10.1037/a0026838

Rosenthal, R., Rosnow, R. L., & Rubin, D. B. (2000). Contrasts and effect sizes in behavioral research: A correlational approach. Cambridge University Press.

Schultz, W. (2016). Dopamine reward prediction-error signalling: A two-component response. Nature Reviews Neuroscience, 17(3), 183–195. 10.1038/nrn.2015.26

Sharma, N. S., Acharya, S. K., Nair, A. P., Matalia, J., Shetty, R., Ghosh, A., & Sethu, S. (2019). Dopamine levels in human tear fluid. Indian Journal of Ophthalmology, 67(1), 38–41. 10.4103/ijo.IJO_568_18

Spring, B., Ockene, J. K., Gidding, S. S., Mozaffarian, D., Moore, S., Rosal, M. C., Brown, M. D., Vafiadis, D. K., & Cohen, D. L. (2020). Better population health through behavior change in adults: A call to action. Circulation, 141(10), e653–e664. 10.1161/CIR.0000000000000732

Webb, T. L., Joseph, J., Yardley, L., & Michie, S. (2010). Using the Internet to promote health behavior change: A systematic review and meta-analysis of the impact of theoretical basis, use of behavior change techniques, and mode of delivery on efficacy. Journal of Medical Internet Research, 12(1), e4. 10.2196/jmir.1376

Wise, R. A. (2004). Dopamine, learning and motivation. Nature Reviews Neuroscience, 5(6), 483–494. 10.1038/nrn1406

